# Glymphatic function in the gyrencephalic brain

**DOI:** 10.1101/2020.11.09.373894

**Authors:** Nicholas Burdon Bèchet, Nagesh C. Shanbhag, Iben Lundgaard

## Abstract

Identification of the perivascular compartment as the point of exchange between cerebrospinal fluid (CSF) and interstitial fluid mediating solute clearance in the brain, named the glymphatic system, has emerged as an important clearance pathway for neurotoxic peptides such as amyloid-beta. However, the foundational science of the glymphatic system is based on rodent studies. Here we investigated whether the glymphatic system exists in a large mammal with a highly gyrified brain. CSF penetration into the brain via perivascular pathways, a hallmark of glymphatic function, was seen throughout the gyrencephalic cortex and subcortical structures, validating the conservation of the glymphatic system in a large mammal. Macroscopic CSF tracer distribution followed the sulci and fissures showing that the gyri enhance CSF dispersion. Three-dimensional renditions from light sheet microscopy showed that CSF influx through perivascular spaces was 4-fold more extensive in the pig brain than in mice. This demonstrates the existence of an advanced solute transport system in the gyrencephalic brain that could be utilised therapeutically for enhancing waste clearance.

## INTRODUCTION

Apart from providing cushioning and buoyancy for the brain, cerebrospinal fluid (CSF) has also been implicated in the maintenance of neural homeostasis and removal of harmful metabolites with several experiments demonstrating communication between the CSF and brain neuropil ^1–3^. The glymphatic hypothesis explains the process of advective CSF-ISF exchange for waste removal from the neuropil to the SAS ^4–6^. The glymphatic system is a brain-wide influx and clearance system formed by a network of perivascular spaces (PVS) that permit the exchange of CSF ^4,5,7^. The advective flow of CSF in the PVS and through the brain parenchyma acts to clear metabolic waste ^4,5,8,9^. This process is dependent on aquaporin-4 (AQP4) water channels which are highly expressed on the astrocyte endfeet that define the outer border of the PVS ^4,10^. The precise mechanisms of glymphatic flux and clearance are as of now incomplete, however, it is known that glymphatic function declines with age and loss of AQP4 while influx is temporally correlated with the systolic phase of the cardiac cycle ^10–12^. Furthermore, glymphatic function is activated in the sleep state and brain states resembling natural sleep in terms prominent low frequency spectra (0-4 Hz), such as under ketamine/xylazine anaesthesia ^12,13^. Platelet-derived growth factor B (PDGF-B), which is necessary for localising pericytes to the vascular wall and thereby contributing to APQ4 anchorage, is required for normal glymphatic development ^14,15^. In terms of manipulations, low doses of alcohol and regular voluntary exercise increase glymphatic function ^16–18^. The relationship between glymphatic function and neurodegeneration has also been widely addressed ^4,13,19,20^. Despite gaps in knowledge, fundamental research has built on the glymphatic hypothesis, forming the building blocks of a new concept of fluid exchange in the brain. Advanced light imaging modalities such as 2-photon, confocal and light sheet microscopy in rodents have demonstrated microscopic glymphatic features including in vivo tracer motion within an AQP4 bounded PVS, and subsequent distribution of tracer to the parenchyma, as well as the entire PVS network of the glymphatic system in whole brains ^4,7,12^. However, the molecular details and knowledge of glymphatic physiology and metabolite clearance stems from rodent studies. In humans, magnetic resonance imaging (MRI) experiments have shown that intrathecally administered gadobutrol moves from the CSF compartment into the brain parenchyma at a rate exceeding that of diffusion ^21–23^. Similar macroscopic observations have also been made in non-human primates after CM injection and MR imaging ^24^. These findings point to the conservation of a glymphatic system from mice to humans, however, the low resolution of MRI is unable to discern a precise route of entry for the tracer. These outcomes highlight that the extent of knowledge on glymphatic physiology in humans compared to rodents is comparatively scarce. Unfortunately, glymphatic studies with the resolution to capture PVS as highways for CSF distribution are highly invasive, severely limiting the potential expansion of knowledge in human subjects. This emphasizes the need for the use of an intermediate species, easily accessible for study and more closely related to humans in order to understand the finer details of glymphatic physiology in large mammals.

To this end, we carried out CSF tracer studies in Landrace pigs, which are comparable in size to humans and have highly gyrified brains of similar macroscopic architecture. Using confocal, electron microscopy (EM) and optical clearing followed by light sheet microscopy, our experiments confirmed an extensive perivascular solute transport far exceeding the levels in mice, and thus is consistent with the existence of a highly developed glymphatic system in gyrified brains of large mammals.

## RESULTS

### Sulci facilitate extensive CSF distribution

Injection of tracers in the CM permits them direct access to the subarachnoid space, which contains CSF and covers the entire surface of the brain. We investigated the macroscopic profile of CSF distribution and whether there were any patterns or favoured paths. The entire cerebral hemisphere of a mammalian brain is supplied by the posterior, middle and anterior cerebral arteries and their branches ^25^. Areas where end branches meet and anastomose are called watershed zones and are typically more vulnerable to cerebral ischaemia ^26–28^. We compared mean tracer intensity in the watershed zone, where the branches of the middle and anterior cerebral artery meet to the whole dorsal surface and found that tracer values in this zone were consistently 50% lower, even in the brains with longer circulation times (mean diff. = 9.097 ± 2.133, p=0.0178, two-tailed paired *t*-test; **Fig. 1 a-c**). This observation made it apparent that the most distal arterial branches might not exhibit the same capacity for perivascular distribution as the major cerebral arteries and their initial branches. Thus, these areas may not only be more vulnerable to ischaemia but could also have a greater propensity for waste aggregation in instances of reduced glymphatic function. Influx differences in the watershed zone did not differ significantly with circulation time so we further addressed this parameter by taking the tracer intensity around the interhemispheric fissure as a maximum intensity value and determined how far laterally tracer travelled before amounting to 50% of the maximal value, here termed “midline shift.” Midline shift was linearly correlated with longer circulation times amounting to a greater shift laterally in tracer intensity, although this correlation was not significant (R^2^=0.9853, p=0.0774, **Fig. 1 a, d**). Thus, with longer circulation times, tracer tends to be transported further laterally from the midline but not all the way to the watershed zone, further reaffirming the role of large calibre arteries and their early branches in driving CSF distribution ^12,23^. Light sheet imaging of a cleared volume of pig cortex additionally highlighted large vessels as a key vehicle in transmitting tracer influx deeper into the brain cortex (**Supplementary video 1**).

**Figure 1.**
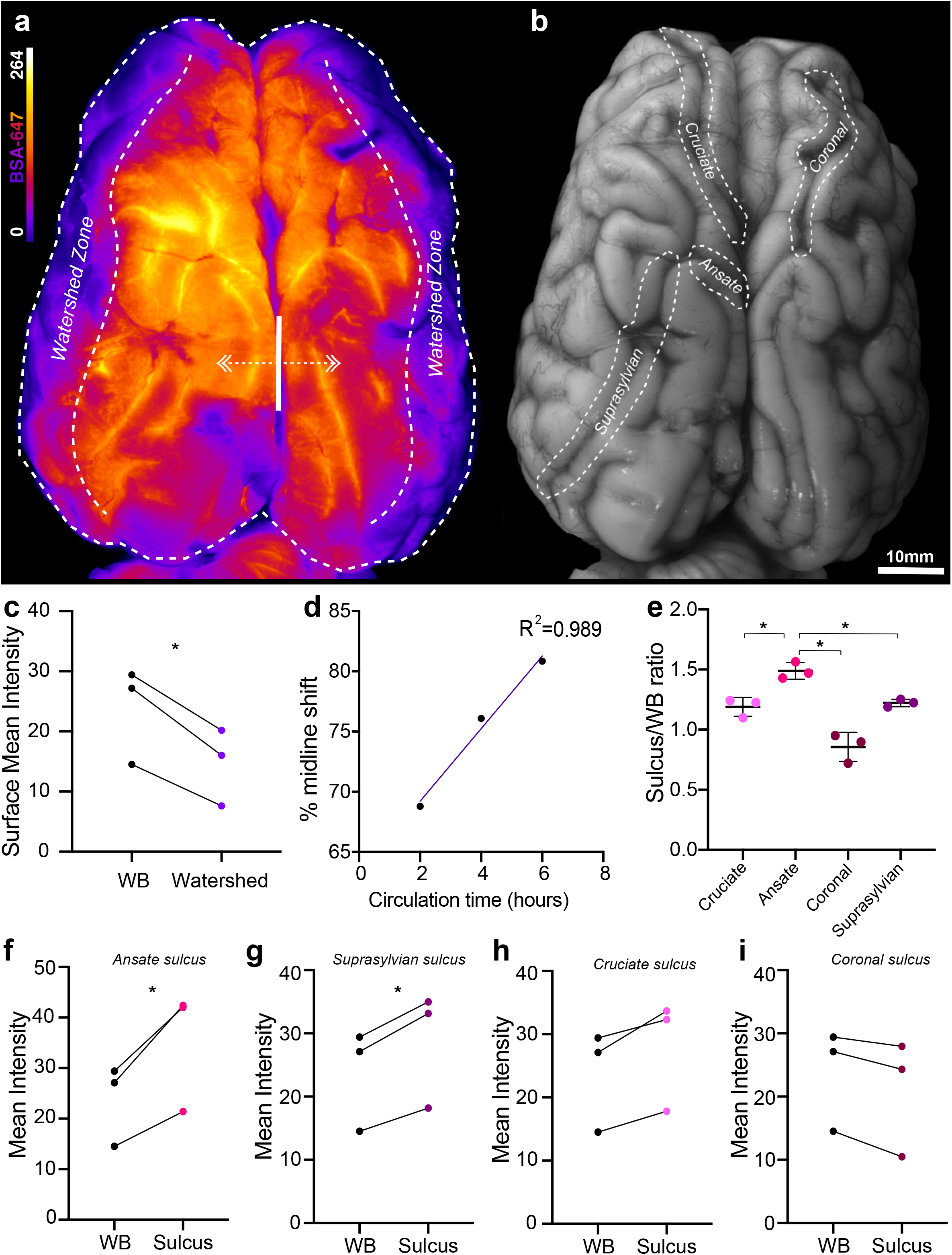
CSF tracer distribution at the dorsal brain surface. **a** Representative image of whole dorsal surface of pig brain under fluorescent light showing BSA-647 tracer distribution. **b** Representative image of whole dorsal surface of pig brain under white light. **c** Surface BSA-647 tracer intensity over whole dorsal surface versus watershed zone (two-tailed paired *t*-test, p=0.0178). **d** Liner regression of “midline shift” vs circulation time. Percentage distance from IHS fissure to lateral aspect of brain where tracer reaches 50% value of IHS value (R^2^=0.989, p=0.0774). **e** Comparison of sulcus/whole brain intensity ratio amongst cruciate, ansate, coronal and suprasylvian sulci (repeated measures one-way ANOVA with Tukey’s multiple comparison). **f** Surface BSA-647 tracer intensity over whole dorsal surface versus ansate sulcus (two-tailed paired *t*-test, p=0.0433). **g** Surface BSA-647 tracer intensity over whole dorsal surface versus suprasylvian sulcus (two-tailed paired *t*-test, p=0.0199). **h** Surface BSA-647 tracer intensity over whole dorsal surface versus cruciate sulcus (two-tailed paired *t*-test, p=0.0674). **i** Surface BSA-647 tracer intensity over whole dorsal surface versus coronal sulcus (two-tailed paired *t*-test, p=0.0671). N=3. *p<0.05. Graphs represent mean ±SD. BSA-647, Alexa Fluor 647 conjugated to bovine serum albumin.

An evolutionary advancement in the brains of higher mammals is the development of sulci and gyri from the folding of the brain ^29^. Sulci increase the tissues’ surface area which our data shows could also impact CSF distribution. On the dorsal surface of the pig brain there are at least 4 major sulci: cruciate, ansate, coronal and suprasylvian ^30^. We determined whether any of the major pig sulci were preferential paths for CSF by comparing the mean tracer intensity surrounding each sulcus at the surface level (**Fig. 1b**). When comparing amongst the sulci, the tracer intensities were significantly higher in the ansate sulcus compared to the cruciate (p=0.0199), coronal (p=0.0391) and suprasylvian (p=0.0477) sulci (repeated measures one way-ANOVA; **Fig. 1e**). When comparing sulci intensities to the entire dorsal surface, both the ansate (mean diff. = 11.6 ± 4.422, p=0.0433, two-tailed paired *t*-test; **Fig. 1f**) and the suprasylvian (mean diff. = 5.097 ± 1.265, p=0.0199, two-tailed paired *t*-test; **Fig. 1g**) sulcus exhibited significantly higher tracer intensities. At the surface level, the cruciate sulcus showed a tendency for higher tracer intensity values (mean diff. = 4.262 ± 2.020, p=0.0674, two-tailed paired *t*-test; **Fig. 1h**) while, interestingly, tracer intensity values tended to be lower in the coronal sulcus when compared to the entire dorsal cerebrum (mean diff. = -2.758 ± 1.304, p=0.0671, two-tailed paired *t*-test; **Fig. 1i**). Both the ansate and cruciate sulci are in open contact with the interhemispheric fissure and thus receive CSF flow directly from this large space, and yet only the ansate yielded significantly higher tracer intensities. This could be because although the cruciate sulcus is in open contact with the interhemispheric (IHS) fissure it is also 4-5 times longer than the ansate, and so the more distal parts of the cruciate with lower intensities bring down its overall intensity. These intriguing findings indicate that brain gyrification is favourable for extensive CSF dispersion throughout the cortical surface.

### Widespread cortical penetration of CSF tracer

To achieve imaging of the full extent of the sulci and their capacity for CSF distribution, we examined two of the large fissures, interhemispheric and rhinal, both at the surface level and from a macroscopic slice level (**Fig. 2 a-c**). The macroscopic slices demonstrated the extent of tracer penetration into both the fissures and sulci (**Fig. 2c**). Compared to the whole lateral surface of the brain, the rhinal fissure yielded significantly higher tracer intensities (mean diff. = 6.322 ± 2.323, p=0.0422, two-tailed paired *t*-test; **Fig. 2d**). This observation was further reinforced at the slice level comparing the fissure intensity to the whole slice, with a mean difference between groups at the slice level more than double that of the surface level (mean diff. = 13.03 ± 2.538, p=0.0002, two-tailed paired *t*-test **Fig. 2e**). The interhemispheric fissure yielded similar results at both surface (mean diff. = 8.415 ± 3.229, p=0.0137, two-tailed paired *t*-test; **Fig. 2f**) and slice levels (mean diff. = 11.96 ± 4.195, p=0.0107, two-tailed paired *t*-test; **Fig. 2g**). This shows that at the given circulation times, more CSF moves through the SAS overlying these regions than other surrounding regions. Similarly in humans, brain regions adjacent to the IHS fissure and Sylvian fissure, comparable to the rhinal fissure in pigs, appear to exhibit earlier and more intense tracer enrichment than surrounding regions ^23^. This could be due to lower fluid resistance derived from more potential space within sulci in combination with arterial pulsations from branches of the large calibre arteries that run within fissures and sulci.

**Figure 2.**
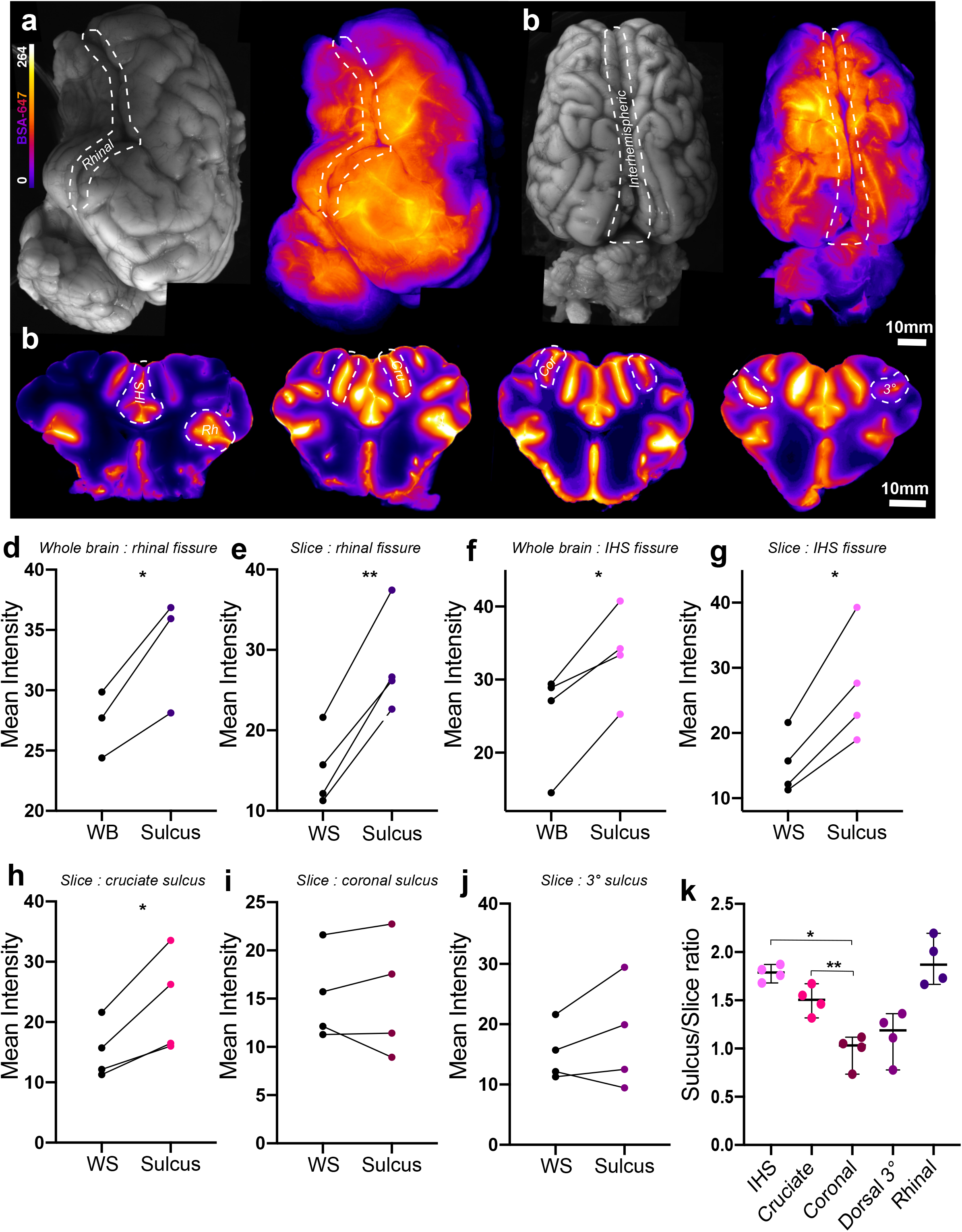
Macroscopic imaging of slices give greater insights into CSF tracer distribution. **a** Representative images of whole lateral surface of pig brain under white light and under fluorescent light showing BSA-647 tracer distribution. **b** Representative images of whole dorsal surface of pig brain under white light and under fluorescent light showing tracer distribution. **c** Representative images of macroscopic brain slices showing tracer distribution. **d** Surface tracer intensity over whole lateral surface versus rhinal fissure. (two-tailed paired *t*-test, p=0.0422, N=3). **e** Tracer intensity over whole slice versus rhinal fissure. (two-tailed paired *t*-test, p=0.002). **f** Surface tracer intensity over whole dorsal surface versus interhemispheric fissure (two-tailed paired *t*-test, p=0.0137). **g** Tracer intensity over whole slice versus interhemispheric fissure. (two-tailed paired *t*-test, p=0.0107). **h** Tracer intensity over whole slice versus cruciate sulcus (two-tailed paired *t*-test, p=0.0281). **i** Tracer intensity over whole slice versus coronal sulcus (two-tailed paired *t*-test, p=0.9794). **j** Tracer intensity over whole slice versus 3° sulcus (two-tailed paired *t*-test, p=0.3219). **k** Comparison of sulcus/whole slice intensity ratio amongst interhemispheric fissure, cruciate sulcus, coronal, 3° sulcus and rhinal fissure (repeated measures one-way ANOVA with Tukey’s multiple comparison). N=4. *p<0.05, **p<0.01, ***p<0.001, ****p<0.0001. Graphs represent mean ±SD. BSA-647, Alexa Fluor 647 conjugated to bovine serum albumin; Cor, Coronal; Cru, Cruciate; HIS, Interhemispheric; Rh, Rhinal; 3°, Tertiary.

The same analyses were carried out on 3 other sulci at the slice level: the cruciate, coronal and so-called “3°” sulcus. Of these only the cruciate sulcus exhibited significantly higher values than the whole slice (mean diff. = 7.881 ± 3.945, p=0.0281, two-tailed paired *t*-test; **Fig. 2h-j**). Comparing tracer intensities between the sulci and fissures revealed that fissures (IHS and rhinal) tended to receive more CSF tracer than the sulci (repeated measures one-way ANOVA; **Fig. 2k**).

Next, tissue bounding the interhemispheric fissure was sectioned and imaged for insights into glymphatic penetration (**Fig. 3a**). Tracer circulation time yielded a linear correlation with both overall slice intensity (R^2^=0.6919, p=0.3746; **Fig. 3b**) and white matter fluorescence intensity (R^2^=0.7743, p=0.3152; **Fig. 3c**), albeit not significant with the number of animals. At this microscopic level we also compared tracer intensities along the slice surface to tracer intensities within the sulci, with values in the sulci consistently higher (mean diff. = 4.995 ± 2.508, p=0.0283, two-tailed paired *t*-test; **Fig. 3d**). This phenomenon was further explored through optically clearing a piece of pig cortex and light sheet imaging and the preference of tracer toward and within a sulcus as compared to the surface was evident (**Supplementary video 2**). Finally, we addressed depth of tracer penetration, a proxy for glymphatic influx, versus tracer concentration and tracer circulation time and observed that longer circulation times yielded trends for both higher tracer surface intensities along with deeper penetration (influx) and more gradual reductions in tracer intensity at regions more distal from the surface (**Fig. 3e**). High magnification epifluorescence and confocal microscopy images further exhibited a dense perivascular network of glymphatic influx in the large mammalian brain (**Fig. 3f-g**).

**Figure 3.**
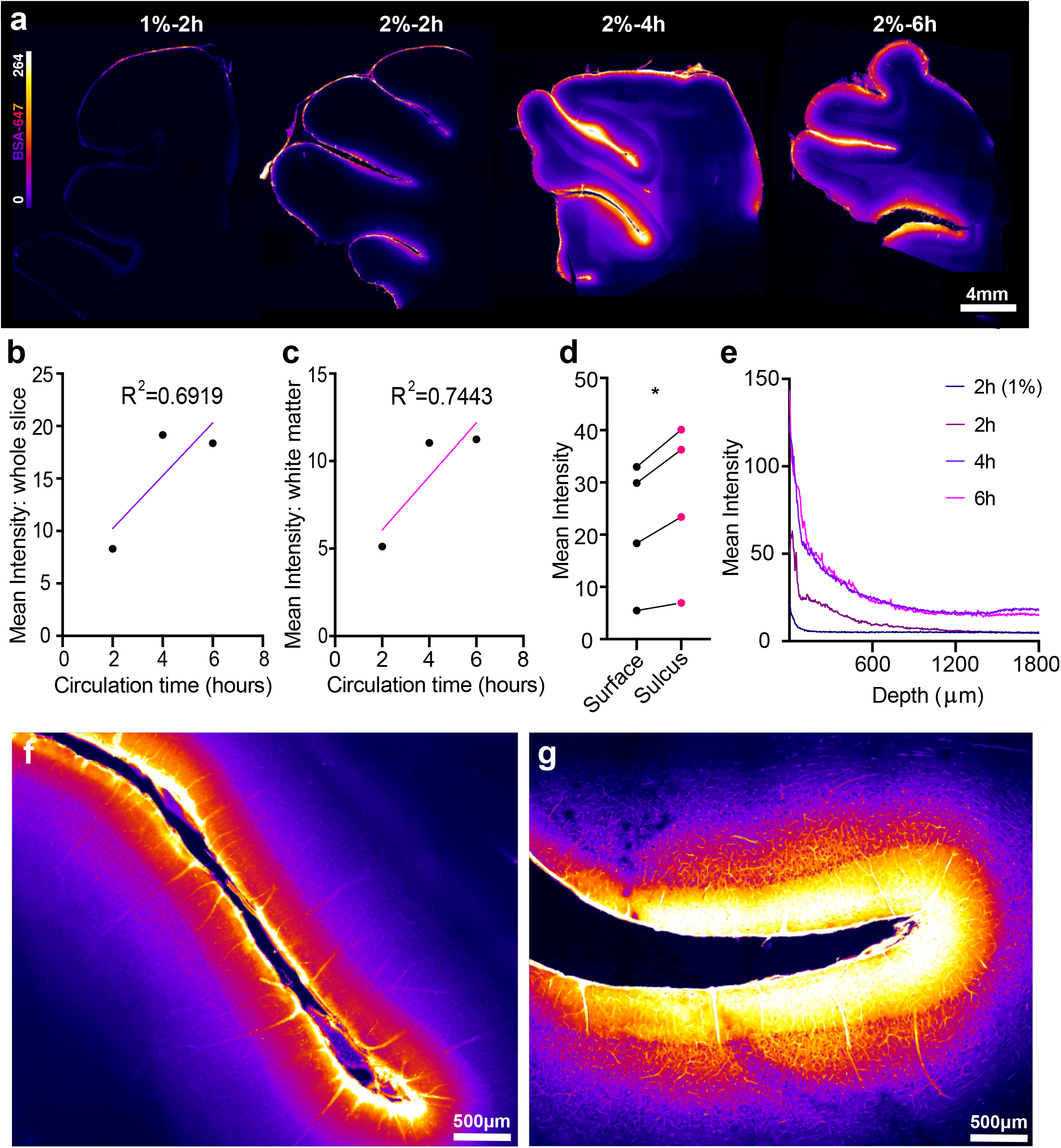
Glymphatic influx predominates at sulci. **a** Representative images of vibratome brain slices demonstrating glymphatic influx. **b** Linear regression of overall slice intensity vs circulation time (R^2^=0.6919, p=0.3764). **c** Linear regression of white matter slice intensity vs circulation time (R^2^=0.7443, p=0.3152). **d** BSA-647 tracer intensity over exposed surface vs tracer intensity over of sulcul surface (two-tailed paired *t*-test, p=0.0283, N=4). **e** Mean pixel intensity from slice surface to 1800 *μ*m into the slice across circulation times and BSA-647 tracer intensities. **f** Representative 20X magnified epiflourescent image of slice sulcus. **g** Representative 20X magnification image of slice sulcus using confocal. *p<0.05. BSA-647, Alexa Fluor 647 conjugated to bovine serum albumin.

The increased tracer intensity in sulci compared to the surface and gyri was clearly visible across imaging modalities which validates the role of these structures in directing CSF, highlighting how the anatomical differences in the gyrencephalic brain as compared to the lissencephalic brain can functionally influence CSF paths.

### CSF penetrates the porcine cortex via perivascular routes

PVS drastically increase the surface area through which the CSF communicates with the brain and is believed to efficiently facilitate the movement of CSF throughout the neuropil and clearance of waste products ^4,12^. The inner bounds of the PVS are formed by endothelial cells while astrocytic endfeet giving rise to the glia-limitans form the outer bound. To visualise tracer in the perivascular spaces of the pig brain, we stained for GLUT-1 (endothelial cells) and AQP4 (**Fig. 4a**). Tracer was evident both along vessels and in the surrounding parenchyma as a haze in upper cortical layers. More distal from the cortical surface, the amount of tracer in the parenchyma lessened while perivascular tracer remained unequivocally present (**Fig. 4b**). Higher magnification confocal images of endothelial cells surrounded on the outer rim by dense AQP4 staining exhibited clearly visible tracer running along the longitudinal aspects of vessels (**Fig. 4c-e**). A complex three-dimensional AQP4 architecture contributing to a bulky perivascular space was seen surrounding vessels of approximately 20 μm in diameter (**Fig. 4d-f**). By plotting an intensity profile for each fluorophore, three distinct peaks demonstrated that CSF tracer is bounded by an endothelial cell and AQP4 peak either side, constituting the PVS (**Fig. 4g**). Similar results emerged when using a GFAP staining for astrocytes (**Fig. 4h-n**) and thus these findings demonstrate the presence of the integral microscopic machinery of the glymphatic system as described in rodents now identified in a large mammal.

**Figure 4.**
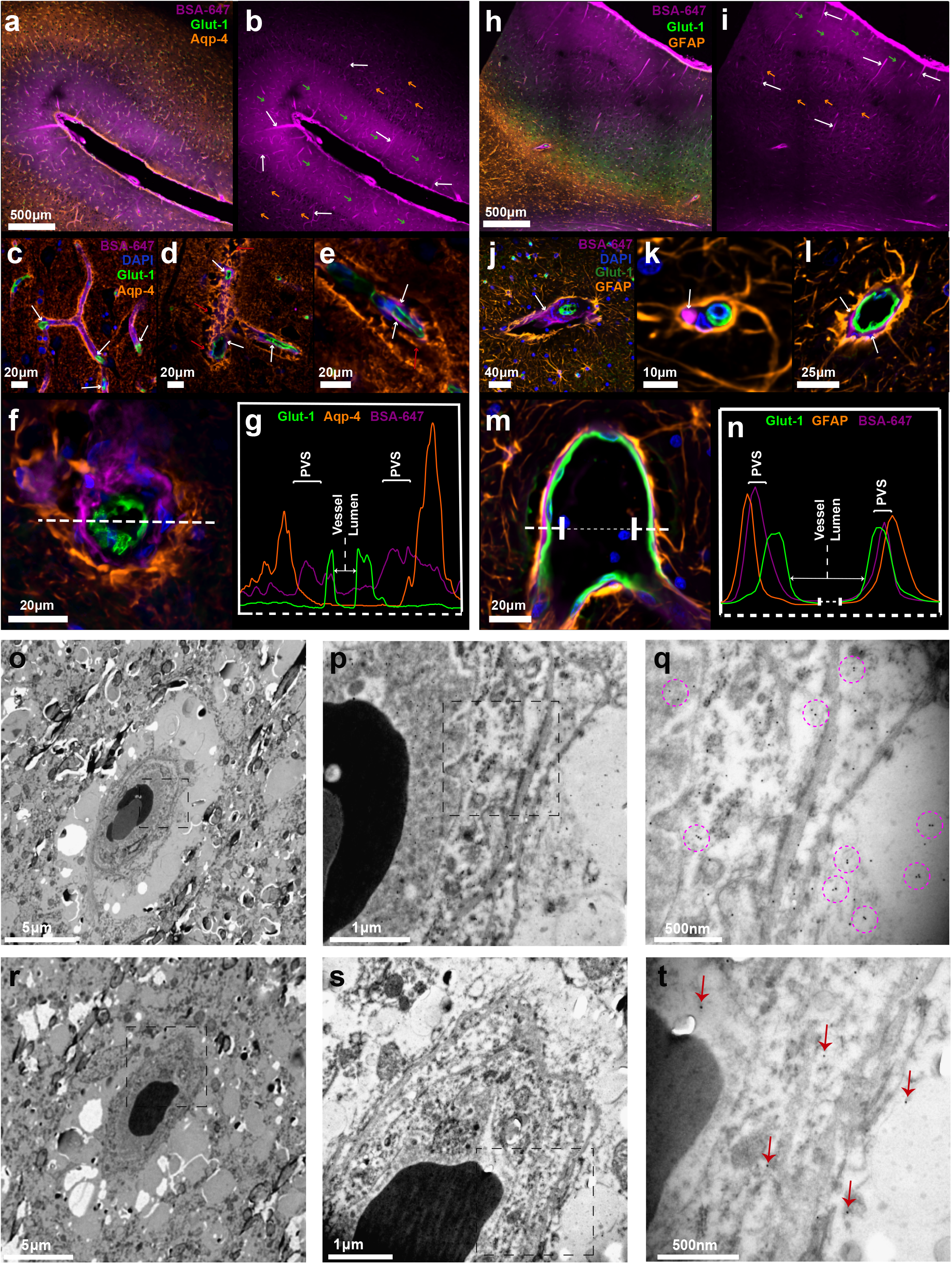
CSF penetrates the porcine cortex via perivascular routes. **a** Representative image of immunohistochemical staining with Glut-1 for endothelial cells and AQP4 with tracer (BSA-647). **b** Representative image of CSF tracer distribution. White arrows point to tracer in perivascular space (PVS). Green arrows show higher amounts of tracer present in the parenchyma near the surface. Orange arrows show lower amounts of tracer visible deeper in the parenchyma. **c-e** 20x magnified images of tracer in the PVS along vessels. White arrows point to perivascular tracer. Red arrows point to AQP4 architecture. **f** Cross section through cortical vessel showing perivascular tracer ensheathed by AQP4. **g** Intensity profiles of AQP4, BSA-647 and Glut-1 across the vessel cross-section. **h** Representative image of immunohistochemical staining with Glut-1 (blood vessels) and GFAP (astrocytes) with tracer (BSA-647). **i** Representative image of tracer distribution. White arrows show tracer in perivascular space (PVS). Green arrows point to higher amounts of tracer present in the parenchyma near the surface. Orange arrows point to lower amounts of tracer visible deeper in the parenchyma. **j-l** 20x magnification images showing tracer in the PVS along vessels. White arrows point to perivascular tracer. **m** Cross section through cortical vessel showing perivascular tracer ensheathed by GFAP. **n** Intensity profiles of GFAP, BSA-647 and Glut-1 across the vessel cross-section. **o-q** Electron microscopy images of a single vessel in pig cortex stained with primary antibodies against tracer and secondary antibodies coupled to gold nanoparticles. Purple circles highlight specific clusters of immunogold staining. **r-t** Electron microscopy images of a single vessel in pig cortex stained with only secondary antibodies coupled to gold nanoparticles as control. Red arrows point to single gold particles. BSA-647, Alexa Fluor 647 conjugated to bovine serum albumin.

To obtain even higher resolution images, we used immunogold labelling against the CSF tracer (bovine serum albumin) and performed electron microscopy (EM). Imaging at 2,000-30,000X on a transmission EM (TEM) platform allowed for the identification of blood vessels and the brain tissue surrounding them (**Fig. 4o-t**). In order to control for non-specific retention of gold nanoparticles we carried out staining with (**Fig. 4o-q**) and without (**Fig. 4r-t**) primary antibodies. While non-specific retention of gold particles in control samples yielded randomly dispersed singular particles (**Fig. 4t**), specific immunogold staining was visible in the experimental group identifiable as several clusters of 2-3 gold nanoparticles in the anatomical region circumventing the vessel (**Fig. 4q**). This further strengthens our findings that tracer moves into the brain along blood vessels and this phenomenon persists down to a capillary level with immunogold staining.

### Subcortical favouritism of hippocampal CSF influx

To investigate subcortical glymphatic influx in the pig, we dissected and sliced the hippocampus from each brain (**Fig. 5a**). Although longer incubation times clearly yielded greater influx in the hippocampus only a weak linear correlation could be modelled (R^2^=0.595, p=0.4392; **Fig. 5b**). In order to understand how glymphatic function is favoured in the hippocampus it was divided into ventral and dorsal regions **(Fig. 5c)**. The ventral aspect of the hippocampus exhibited significantly more glymphatic influx than the dorsal region (mean diff. = 0.786 ± 0.1839, p=0.0034, two-tailed paired *t*-test; **Fig. 5d**). Both the dentate gyrus (DG) and the entorhinal cortex (ERC) exhibited significantly more tracer influx than in CA1 and CA2, which were both comparable (repeated measures one-way ANOVA; **Fig. 5e-f**). These findings can be resolved in an anatomical sense as the ventral aspect of the hippocampus, and thus the ventral regions like the DG and ERC, are in direct communication with the ambient cistern. Thus, CSF and tracer have more direct and immediate access to the hippocampus.

**Figure 5.**
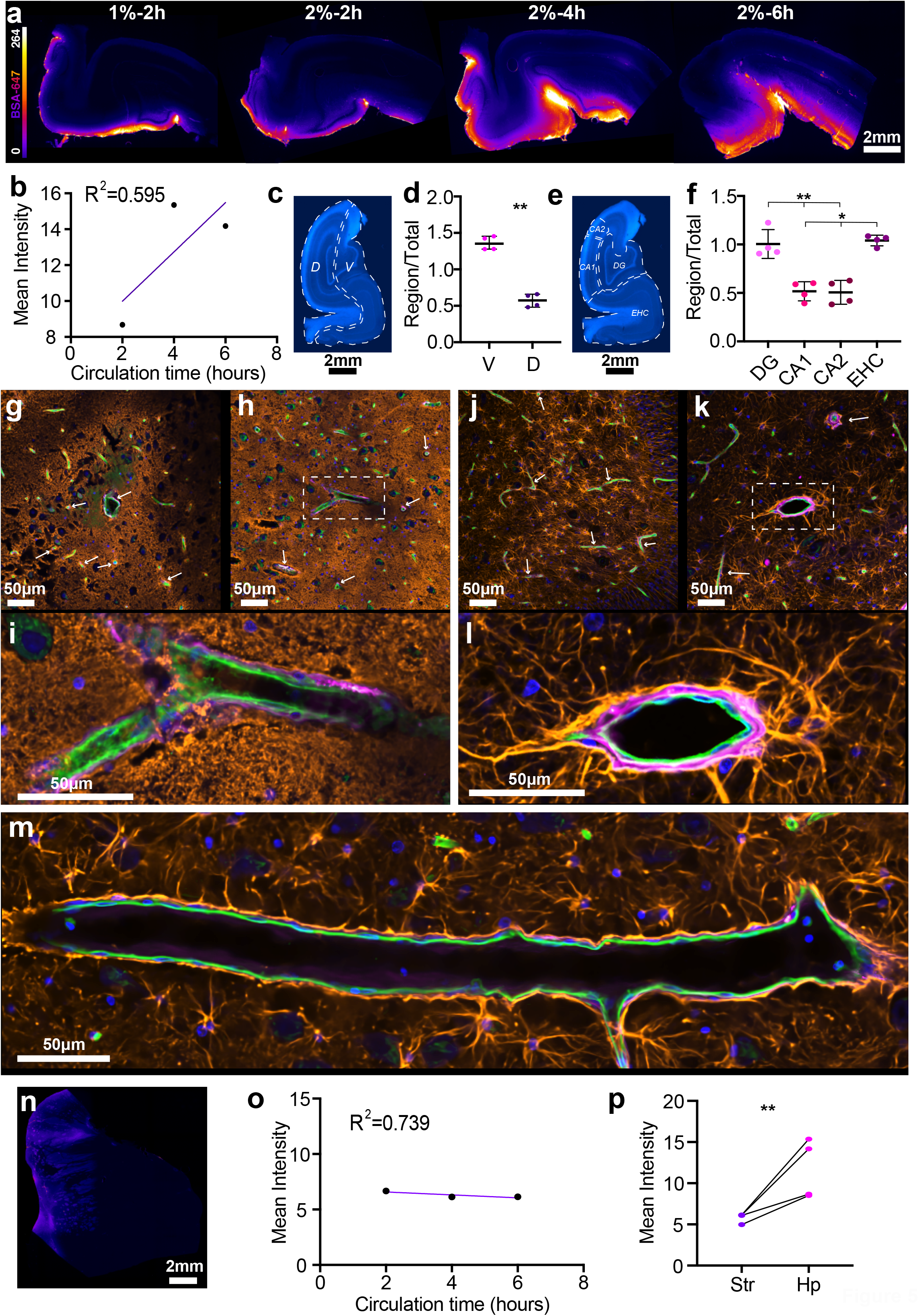
Glymphatic influx reaches hippocampus before striatum. **a** Representative images of vibratome hippocampal slices showing glymphatic influx. **b** Linear regression of overall hippocampal intensity vs circulation time (R^2^=0.595, p=0.4392). **c** Representative image of pig hippocampus stained with DAPI showing dorsal/ventral division. **d** Comparison of regional vs whole slice intensity ratios for dorsal and ventral hippocampal regions (two-tailed paired *t*-test, p=0.0034). **e** Representative image of pig hippocampus stained with DAPI showing regional divisions. **f** Comparison of regional vs whole slice intensity ratios for CA1, CA2, entorhinal cortex and dentated gyrus (repeated measures one-way ANOVA with Tukey’s multiple comparison). **g-i** Representative images of hippocampus stained with AQP4, lectin and DAPI with BSA-647 tracer. White arrows point to tracer in PVS. **j-l** Representative images of hippocampus stained with GFAP, lectin and DAPI with tracer. White arrows point to tracer in PVS. **m** 20x magnified image of hippocampus showing longitudinal-sectional face of blood vessel stained with lectin with tracer in the PVS and GFAP staining for astrocytes. **n** Representative image of pig striatum. **o** Linear regression of overall striatal intensity vs circulation time (R^2^=0.739, p=0.3416). **p** Comparison of mean striatal vs mean hippocampal intensity (two-tailed paired *t*-test, p=0.0376). N=4. **p<0.01, ***p<0.001. Graphs represent mean ±SD. BSA-647, Alexa Fluor 647 conjugated to bovine serum albumin.

Vascular staining in combination with AQP4 or GFAP showed the network of astrocytes in the hippocampus. GFAP immunoreactivity revealed that astrocyte endfeet visibly projected to the PVS and CSF tracer was visible within the AQP4 lining surrounding hippocampal vessels (**Fig. 5g-m**).

Interestingly, in contrast to glymphatic influx in the hippocampus, which showed a trend to increase with longer circulation times, influx into the striatum did not differ between 2, 4 or 6 hours (R^2^=0.7386, p=0.3416; **Fig. 5n-o**). Furthermore, overall CSF tracer influx was significantly higher in the hippocampus compared to the striatum (mean diff. = 5.842 ± 3.274, p=0.0376, two-tailed paired *t*-test; **Fig. 5p**). This is in keeping with a previous study in humans, where hippocampal tracer enrichment began to increase after 2 hours finally peaking around 8 hours and tracer enrichment in the basal ganglia peaked at only 48 hours ^23^.

### Perivascular CSF transport in pigs is more developed than in rodents

Light sheet microscopy of optically cleared tissues enables imaging of whole tissue volumes at optical resolution without the need for tissue sectioning. Recently, our lab showed that visualisation of the glymphatic system in entire mouse brains is possible using optical tissue clearing and light sheet microscopy and provides a full overview of glymphatic architecture ^7^. To compare the glymphatic system in pigs and mice in undisturbed brain tissue, we applied optical clearing and light sheet microscopy which revealed a pattern of regularly distributed entry points of CSF tracer from the cortical surface into the brain (**Fig. 6a-b**). Quantification of the influx density of CSF tracer per millimetre of cortical surface revealed a robust 4-fold greater influx into perivascular spaces in pigs compared to mice (39.33 ± 1.607 vs 10.5 ± 3 PVS/mm^2^, p<0.0001, students *t*-test; **Fig. 6c-e**). This further translated to a superior area-coverage of cortical surface by perivascular influx (4.497 ± 0.8648 vs 2.037 ± 0.1897 % area coverage, p=0.0086, students *t*-test; **Fig. 6f**). The vast extent of CSF distribution to perivascular spaces in the pig brain was further visible in three dimensional renderings of the cortical surface (**Supplementary video 3-4**).

**Figure 6.**
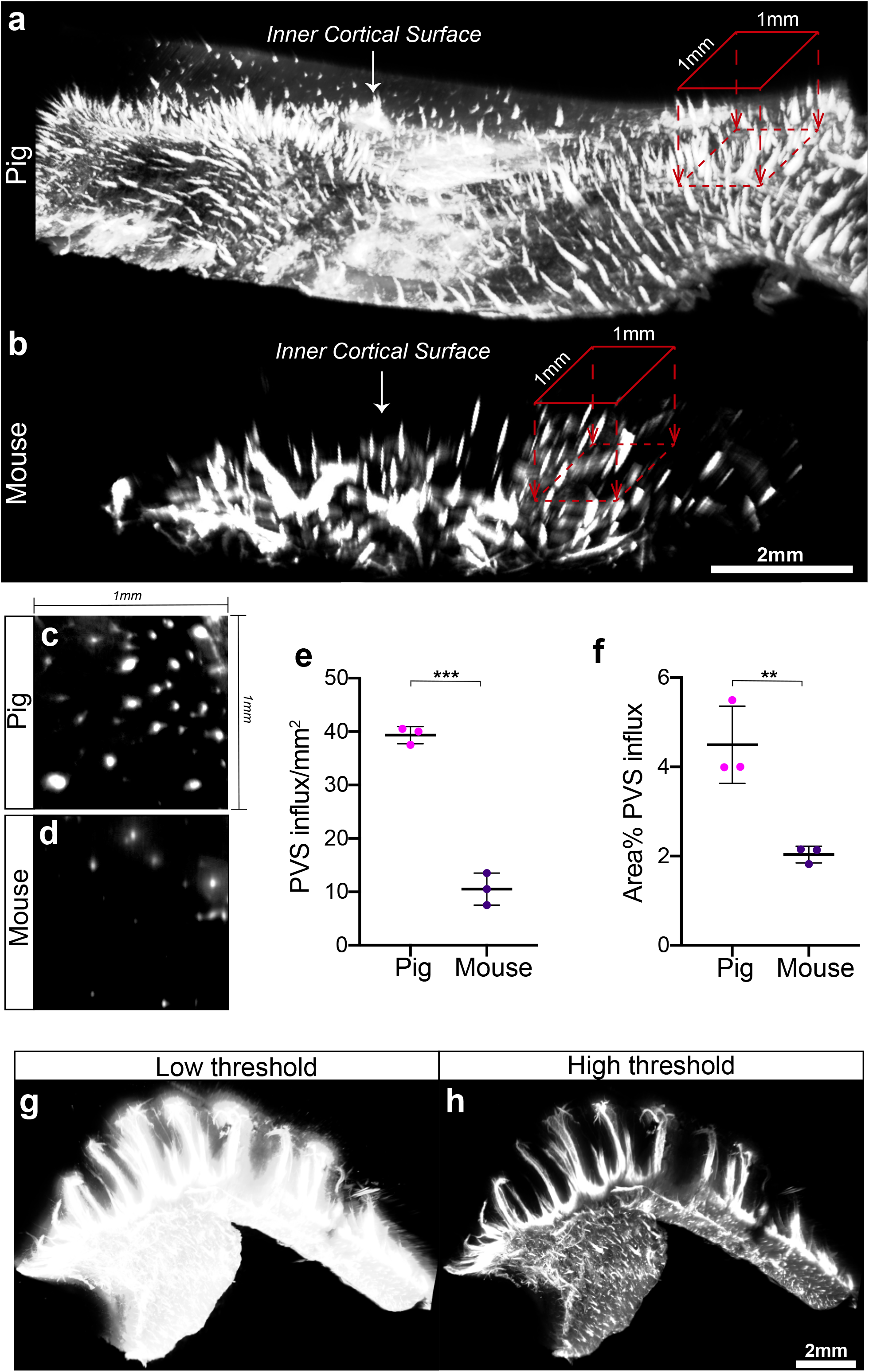
The extent of CSF tracer distribution in the pig PVS network exceeds that in mice. **a-b** 3D reconstruction of pig and mouse inner cortical surface highlighting perivascular influx sites. 1mm by 1mm region of interest is used to count perivascular influx sites projecting into the brain. Scale bar = 2mm. **c-d** Representative image of a cross section through perivascular channels parallel to cortical surface in pig brain and mouse brain. **e-f** Quantification of number of perivascular influx sites per mm^2^ of cortical surface in pig and mouse brains (students *t*-test, p<0.0001) and percent of cortical surface area covered by perivascular influx in pig and mouse brains (students *t*-test, p=0.0086). **g-h** 3D renditions of pig hippocampus from light sheet microscope at low and high thresholds. N=3. **p<0.01, ***p<0.0001. Graphs represent mean ±SD.

To explore CSF tracer influx in a region with a specialised tissue architecture, hippocampi were optically cleared and light sheet imaged. A low threshold reconstruction exhibited pervasive CSF tracer influx into the entire hippocampal volume while a high threshold visualised the primary pathways which produced this pervasive influx (**Fig. 6g-h**). These primary pathways for CSF influx can be further appreciated in 3D renderings of the entire sub-cortical volume (**Supplementary video 5-6**).

In summary, optical clearing and light sheet microscopy revealed a highly organised pattern of CSF entry into the brain across species and that CSF transport is enhanced in the gyrencephalic brain.

## DISCUSSION

The recently discovered glymphatic system is a brain clearance pathway that has been implicated in the removal of toxic peptides such as Aβ, which is strongly linked to the pathogenesis of AD. Perivascular spaces represent a key constituent of the glymphatic system, providing an extensive network for CSF-ISF exchange ^4,5^. The vast majority of all investigations into the glymphatic system and our most fundamental knowledge of it stems from rodent studies ^4,10,11,13,15,16,31,32^. The few studies carried out in humans utilized MRI and were valuable in demonstrating macroscopic cortical influx patterns, yet lacked the spatial resolution to demonstrate the microscopic glymphatic machinery such as perivascular influx and polarised expression of AQP4 ^21–23,33^. Similar macroscopic contrast enhancement was also revealed in macaques after CM injection with gadolinium chelate ^24^. Herein, we demonstrated the extent of the glymphatic system in a gyrified brain from a large mammal, ranging from specific patterns of CSF paths at the whole brain level, thought to be mediated by a folded architecture in combination with arterial pulsations from large calibre blood vessels, down to the perivascular localisation of CM-injected tracers in both the cortex and subcortical structures. Furthermore, we have used light sheet imaging of cleared brain volumes to visualise the three-dimensional glymphatic architecture of the gyrencephalic brain at optical resolution which enabled a direct comparison of the extent of the PVS network across species. This data showed that perivascular-mediated influx of tracer into the brain, identified as one of the key premises for glymphatic function, is up to 4 times greater in the pig brain than in the mouse brain.

Animal models form a central part of scientific inquiry concerning basic physiology and disease ^34–36^. Rodents serve as the primary model of choice for experimentation and were pivotal in the discovery and characterization of the glymphatic system. However, rodents differ substantially from humans, and the existence and physiology of the glymphatic system needs to be unravelled in the large gyrencephalic brain ^37–39^. The mouse brain weighs around 0.5 g and is lissencephalic while a human brain approximately 3000 times larger and is gyrified ^40,41^. Mice are nocturnal and sleep for multiple short periods throughout the day while most humans require an average of 8 hours of single extended bouts of sleep to maintain cognitive health ^42,43^. When investigating the glymphatic system where size and distance are a factor for convective and/or diffusive CSF-ISF exchange and where sleep and sleeping behaviours strongly govern its function, it becomes more important to work in animal models that more closely match these variables in humans. Interestingly, the pig brain is both closer in size and more similar in macroscopic structure to a human brain than that of a rhesus macaque ^44^. Pigs also appear to be among the mammals with a sleeping pattern that most closely resembles that of humans ^45^. While humans sleep approximately 33.3 % of 24 hours, pigs are documented to sleep for approximately 32.6 % of this period, with the rhesus macaque coming in at 49.2 % and mice on average sleeping for 44% of this period, but highly fragmented ^45^. The pig is therefore a suitable intermediate species between mice and humans for the field of glymphatics and pushes the frontiers of our understanding of the glymphatic system and translating this understanding to humans.

A primary concern when introducing exogenous tracers into the CSF is raised intracranial pressure (ICP). An increase in the volume of any cranial constituent will transiently raise the ICP until that constituent is normalised or is compensated for ^46^. Thus, we acknowledge that the intrathecal administration of tracers into the CSF could raise the ICP, however at low volumes this elevation is momentary with normalisation to baseline in minutes ^47^. Furthermore, if the injection is given at low volumes over time as opposed to a bolus dose ICP is not significantly elevated, as was the case at a rate of 1.6 μL/min infusion in rats ^47^. In mice, it has been accepted to inject 10 μL with an infusion rate of 1 μL/min for glymphatic studies ^7,10^. Translating this to the pigs, whose approximate brain mass is 100 g, it would be comparable to inject tracer at a rate of 200 μL/min amounting a final volume of 2 mL. Instead, we injected ¼ of the translated acceptable volume at ½ the infusion rate and it is thus unlikely that maintained pathological elevations in ICP occurred. Thus, it is unlikely that our observations reflect results from aberrations in ICP.

A key debate amongst researchers in the field of CSF dynamics and the glymphatic system is the nature of CSF-ISF exchange, concerning whether it be by diffusion, convection, or a combination of both, advection. Some arguments against the glymphatic system are based on modelling and state that ISF-CSF exchange could be explained by diffusion alone ^48,49^. Although it is known that sleep further expands the extracellular space which helps drive convective glymphatic influx and clearance, the pressure required to overcome the large calculated hydraulic resistance of the rodent 20-70 nm extracellular space is a point of criticism ^13,48,50–53^. However, the early work from Cserr and colleagues showed that molecules with up to a 5-fold difference in diffusion coefficient exhibited highly similar clearance rates, which should not be the case if diffusion were the main driver ^54^. With mounting evidence from in vivo imaging of tracers and nanoparticles it is now more accepted that the motion of CSF, at least in pial perivascular spaces, is convective in nature and largely driven by arterial pulsations ^4,12^. Pioneering human studies using intrathecal injections of gadobutrol show a degree of contrast enhancement exceeding that predicted by diffusion modelling ^23^. In another follow up study in humans, a more complex model accounting for the geometry of the brain along with several other parameters was applied and once again the degree of observed enhancement exceeded that predicted based on diffusion tensor imaging ^33^. Our data resolves why the tracer circulation in a gyrencephalic brain is faster than diffusion given the marked 4-fold increase in the number of perivascular spaces per unit surface area there is a 4 times greater capacity for convective transport of tracers into and out of the brain as well as a reduction is distances needed for clearance of molecules. Thus, this architectural phenomenon may account for the rapid contrast enhancement seen in human studies and gives insights into glymphatic physiology in the large mammalian brain not seen before. This study negates that the glymphatic system is a rodent-specific phenomenon and strengthens the notion of bulk flow in the perivascular space as a means of overcoming large distances for solute exchange.

CSF-based clearance could account for 25% of Aβ clearance from the mouse brain suggesting that glymphatic function is a significant player in maintaining Aβ homeostasis ^55,56^. Expanding links have been made in rodents between ageing and impaired glymphatic function and this fits with the hypothesis that lack of glymphatic function contributes to neurodegeneration ^11,20^. Knock out of AQP4, which is the main molecular machinery identified as necessary for normal glymphatic function, reduces the clearance rate of Aβ to a third compared to wild type ^4^. Knock out of AQP4 in APP/PS1 mice exacerbates AD plaque formation and significantly enhances Aβ plaque deposition ^20^. Taken together, these findings are encouraging in terms of potentially utilizing glymphatic function as a therapeutic target against neurodegeneration. The only supportive evidence collected from humans is a post mortem study which revealed that AQP4 polarisation had a strong association with AD status and pathology ^57^. As such the potential role of glymphatic function in neurodegeneration would benefit from being further explored in large mammals with a brain similar to that of humans. Through this research we have taken a stride forward in uncovering what might closely resemble the CSF-ISF exchange that takes place in the human brain, and thereby shed light on the more likely architecture of human glymphatics. In this way, our work serves as a base to progress in validating findings from rodents and in doing so maximise the possibility of an eventual effective translation in glymphatic therapeutics to humans.

## METHODS

### Animals

Adult male pigs, *Sus scrofa domesticus*, weighing 50-55 kg, and adult male C57BL/6 mice were used for the experiments. Mice were housed in standard laboratory conditions with a 12h light-dark cycle, *ad libitum* access to water. Pigs were housed in two’s in pens with a 12h light-dark cycle, *ad libitum* access to water. All experimental procedures were performed according to ethical approval by the Malmö-Lund ethical Committee on Animal Research (Dnr 5.2.18-10992/18 and Dnr 5.8.18-08269/2019) and conducted according to the CODEX guidelines by the Swedish Research Council, Directive 2010/63/EU of the European Parliament on the protection of animals used for scientific purposes and Regulation (EU) 2019/1010 on the alignment of reporting obligations. This study complies with the ARRIVE (Animal Research: Reporting in Vivo Experiments) guidelines for reporting of animal experiments.

### Anaesthesia

Pigs were first tranquilised/premedicated with an intramuscular injection of Zoletil (tiletamine 3.75 mg/kg + zolazepam 3.75 mg/kg) and Dexdomitor (dexmedetomidine 37.5 μg/kg). Once unconscious, animals were intubated and a 20G cannula was inserted into the ear vein. For maintenance anaesthesia a triple-drip (100 ml ketamine 100 mg/ml (5 mg/kg/min), 200ml fentanyl 50 μg/ml (2.5 μg/kg/min), 100 ml midazolam 5mg/ml (0.25 mg/kg/min) was applied through the ear vein until effect (± 0.5ml/10kg/min). Breathing was maintained at 14 breaths per minute using a Servo ventilator. All animals underwent constant monitoring for heart rate, blood pressure, oxygen saturation, pO_2_, pCO_2_ and subsequent anaesthesia infusion rate adjustment if needed. Mice received a single intraperitoneal injection of ketamine (100 mg/kg) and xylazine (20 mg/kg).

### Exposure of cisterna magna

Briefly, the skin overlying the back of the head and neck was resected. The underlying muscle layers were severed at their respective origins and retracted. Any excess tissue overlying the skull base and atlas was removed.

### Intracisternal tracer infusion

For pigs while the head of the animal was flexed an 18G cannula was introduced approximately 5mm into the cisterna magna and fixed in place with glue and dental cement. 500 μL of either 1 % or 2 % AlexaFluor647-conjugated bovine serum albumin (BSA-647, Invitrogen) was injected using a 1 ml syringe connected to a 10 cm I.V line at a rate of 100 μL per minute. After injection, BSA-647 was allowed to circulate for 2, 4 or 6 hours. For mice, CM injection was carried out with a 30G dental needle (Carpule, Sopira) connected to a 100 µL Hamilton syringe via PE10 tubing. 10 µL of 2 % AlexaFluor647-conjugated bovine serum albumin (BSA-647, Invitrogen) tracer were injected into the CM at 1 µL/min using an KDS Legato 100 single infusion syringe pump. After injection, BSA-647 was allowed to circulate for 30 minutes.

### Tissue processing

Whole brains were carefully extracted by removing the dorsal skull surface with a hand-held rotating saw blade (Dremel, DSM/20) and severing the spinal cord, pituitary gland and cranial nerves with a surgical spatula. Whole brains were post fixed in 4 % paraformaldehyde (PFA) for 24 hours. Brains were then sliced coronally using a salmon knife and slices were fixed in PFA for a further 24 h. For immunohistochemistry and microscopic investigations parts of the brain were sliced with a vibratome (Leica VT1200S) at a thickness of either 100 μm for hippocampus and striatum or 200 μm for the cortex.

### Optical tissue clearing

The iDISCO+ protocol was carried out as explained by Renier *et al* (2016). Pig brain pieces and whole mouse brains were dehydrated in increasing methanol/H_2_O series (20%, 40%, 60%, 80%, 100%, 100%, 1 hour each), delipidated with methanol/dichloromethane (33%/66% for 3 hours) and pure dichloromethane (2 x 15 min), and optically cleared by impregnation dibenzyl ether (DBE) for at least 14 days prior to imaging.

### Immunohistochemistry

Free-floating brain sections were permeabilized and blocked for 45 min at 4 °C in a solution of 1 % BSA, 0.5 % Triton X-100 and 5 % normal donkey serum in PBS. Primary antibodies (rabbit anti-AQP4, 1:500, MerckMillipore; rabbit anti-GFAP, 1:500, Agilent Technologies; mouse anti-GLUT1, 1:250, Abcam) were added in PBS and incubated overnight at 4 °C on a rocking table. After 3×10 min washes in PBS at room temperature secondary antibodies (Alexa-Fluor 488- and 568-conjugated secondary antibodies, 1:1000) in PBS were added for 90 minutes at 4°C on a rocking table. Slices were then washed in PBS with DAPI (1:1000) and/or tomato lectin (Lycoperiscon esculentum, 1:20, SigmaAldrich) for 20 minutes, washed again and mounted.

### Imaging

Whole brains and macroscopic slices were imaged using a Nikon SMZ25 stereomicroscope with a Plan Apo 0.5x objective (0.08 NA) equipped with an Andor Zyla 4.2 Plus sCMOS camera (Mag-0.75x, Zoom-1.5x). The excitation wavelength was 635 nm using a CoolLED pE4000 LED illumination and the emission filter used was a quadruple bandpass filter.

Vibratome slices were imaged with both Nikon Ti2 Eclipse and Nikon A1RHD confocal microscopes. Cleared pig brain tissue and whole cleared mouse brains were imaged using an Ultramicroscope II light-sheet microscope (LaVision Biotech) with a 1.3X LaVision LVMI-Fluor lens (0.105 NA) equipped with an sCMOS camera (Andor Neo, model 5.5-CL3). The excitation wavelength was 640 nm and the emission filter used was 680/30 nm. Brain pieces were imaged immersed in DBE in the transverse orientation at a z-step size of 5 μm with ImspectorPro64 (LaVision Biotec).

### Analyses

For all linear regressions, a single data point for the respective measurement from each pig that received a 2 % tracer injection was plotted against time (2, 4 or 6 hours). For all paired measurements, the mean intensity of a specific region (sulcus, fissure, brain region) was compared to the mean intensity of the whole area of interest (dorsal brain surface, lateral brain surface, whole slice) in the same brain. For independent comparisons, ratios of a specific region of interest (sulcus, fissure, brain region) divided by the mean intensity of the whole area of interest (dorsal brain surface, lateral brain surface, whole slice) in the same brain were used. Pig 3 was excluded from sulcul surface analyses owing to a surface congenital malformation. For “midline shift”, a line was drawn from and perpendicular to the interhemispheric fissure in the posterior cerebral cortex out the lateral brain border; the intensity around the IHS fissure was used as a maximum value and the distance in % from the IHS fissure to the lateral border for the tracer intensity to reach 50 % of the maximum value was used; an average was generated between both cortices in a single brain. For vessel peak plots, a line was drawn through the vessel in an image stack containing each staining; a plot was generated for each fluorophore and all plots were then overlaid on a single graph. For PVS counting, orthogonal views to the inner cortical surface were generated to gain a cross-sectional view of the spaces. Briefly, background was reduced, signal threshold applied, then converted to binary using a mask and particles counted to generate PVS number per mm^2^ and % area coverage. All images were analysed using Fiji ^58^.

### Immunoelectron microscopy

Pig cortical tissues (1 mm x 2 mm) were pre-fixed in a solution containing 1 % PFA and 1.5 % glutaraldehyde in 0.1 M phosphate buffered saline for 3 h at room temperature and then rinsed several times prior to fixation with 1 % osmium tetroxide. This was followed by dehydration with acetone (30-100 %), impregnation and embedding in pure Epon for sectioning (60 nm). Immunostaining was performed using primary antibody against BSA (1:1000, Sigma Aldrich) and secondary antibody conjugated to 10 nm gold nanospheres (1:20, Abcam). Sections were stained with 4 % uranyl acetate for 20 mins followed by 0.5 % lead citrate for 2 minutes to increase the contrast of the tissue. Sections were examined using FEI Tecnai Biotwin 120kv transmission electron microscope (TEM) and photographed using Olympus Veleta 2×2k camera at magnifications ranging from 2000-30,000×.

### Statistics

All statistics were performed on GraphPad Prism 8 (GraphPad Software). Data were tested for normality using Shapiro-Wilk test. Two-tailed paired t-tests or two-tailed students t-tests were used for comparing two groups. Multiple groups were analysed using a one-way ANOVA with Tukey’s multiple comparison post hoc test. All values are expressed as mean ± SD. N represents number of biological replicates. P<0.05 was accepted as statistically significant.

### Data availability

Data supporting the findings from this study are available from the authors upon request.

## Supporting information

Supplementary video 1: Light sheet reconstruction of pig cortex exhibiting tracer influx along the PVS of a large calibre vessel.

Supplementary video 2: Light sheet reconstruction of pig cortex exhibiting more intense tracer distribution in sulcus than cortical surface.

Supplementary video 3: Inner pig cortex demonstrating extensive influx of tracer into brain via regularly distributed perivascular channels.

Supplementary video 4: inner mouse cortex demonstrating influx of tracer into brain via perivascular channels but less than in pig.

Supplementary video 5: Whole pig hippocampus exhibiting primary tracer influx paths from ventral aspect and along large calibre vessels.

Supplementary video 6: Whole pig hippocampus exhibiting primary tracer influx paths from ventral aspect and along large calibre vessels.

## ACKNOWLEDGEMENTS

This work was funded by the Knut and Alice Wallenberg Foundation, Hjärnfonden, the Crafoord Foundation, Vetenskapsrådet (Dnr 2018-02340), and a postdoctoral stipend for NS from the Wenner-Gren Foundations. Lund University Bioimaging Centre (LBIC) is gratefully acknowledged for providing access to Nikon confocal system A1RHD, FEI Tecnai Biotwin 120kv transmission electron microscope and Arrivis analysis platform. We would also like to thank Dr. Malin Parmar, Dr. Anders Björklund and MultiPark for access to the LaVision II Ultramicroscope platform at Lund University Medical Faculty and Bengt Mattsson for expert technical assistance.

## AUTHOR CONTRIBUTIONS

NBB, NCB and IL designed the experiments, NBB and NCB performed the experiments, NBB and IL analysed the data and made figures, NBB, NCB and IL wrote the manuscript, IL contributed with reagents.

## MATERIALS & CORRESPONDENCE

Iben Lundgaard

## SI APPENDIX

**Supplementary video 1:** Light sheet reconstruction of pig cortex exhibiting tracer influx along the PVS of a large calibre vessel.

**Supplementary video 2:** Light sheet reconstruction of pig cortex exhibiting more intense tracer distribution in sulcus than cortical surface.

**Supplementary video 3:** Light sheet reconstruction of inner pig cortex demonstrating extensive influx of tracer into brain via regularly distributed perivascular channels.

**Supplementary video 4:** Light sheet reconstruction of inner mouse cortex demonstrating influx of tracer into brain via regularly distributed perivascular channels but less than in pig.

**Supplementary videos 5-6:** Light sheet reconstruction of whole pig hippocampi exhibiting primary tracer influx paths from ventral aspect and along large calibre vessels.

